# Anabolic SIRT4 exerts retrograde control over TORC1 signalling by glutamine sparing in the mitochondria

**DOI:** 10.1101/635565

**Authors:** Eisha Shaw, Manasi Talwadekar, Nitya Mohan, Aishwarya Acharya, Ullas Kolthur-Seetharam

## Abstract

Anabolic and catabolic signalling mediated via mTOR and AMPK have to be intrinsically coupled to mitochondrial functions for maintaining homeostasis and mitigate cellular/organismal stress. Although, glutamine is known to activate mTOR, if/how differential mitochondrial utilization of glutamine impinges on mTOR signalling is less explored. Mitochondrial SIRT4, which unlike other sirtuins is induced in a fed state, is known to inhibit catabolic signalling/pathways through AMPK-PGC1a/SIRT1-PPARa axis and negatively regulate glutamine metabolism via TCA cycle. However, physiological significance of SIRT4 functions during a fed state is still unknown. Here, we establish SIRT4 as key anabolic factor that activates TORC1 signalling and regulates lipogenesis, autophagy and cell proliferation. Mechanistically, we demonstrate that the ability of SIRT4 to inhibit anaplerotic conversion of glutamine to α-ketoglutarate potentiates TORC1. Interestingly, we also show that mitochondrial glutamine sparing or utilization is critical for differentially regulating TORC1 under fed and fasted conditions. Moreover, we conclusively show that differential expression of SIRT4 during fed and fasted states is vital for coupling mitochondrial energetics and glutamine utilization with anabolic pathways. These significant findings also illustrate that SIRT4 integrates nutrient inputs with mitochondrial retrograde signals to maintain a balance between anabolic and catabolic pathways.

## Introduction

It is intuitive that the ability to uptake and utilize macronutrients for catabolic or anabolic purposes is intrinsically linked with cellular and organismal energetics. Thus, the sensing and utilization of various macronutrients in the mitochondria needs to be coupled to the activity of metabolic sensors in the cytosol such as AMPK (AMP-activated kinase) and mTOR (mammalian Target of Rapamycin), in order to orchestrate a balance between the metabolic state of the cell and external stimuli. AMPK, mTOR and Sirtuins (Sir2-like NAD-dependent deacylases) have been well established to play central roles in linking nutrient and energetic status to cellular and organismal physiology (1–5). While the AMPK-Sirtuin (6–9) and AMPK-mTOR (10, 11) cross talks are well worked out, the relative interdependence and hierarchy of signals between these sensors is not well explored. Further, functional interactions between Sirtuins and mTOR are poorly understood, and are largely limited to SIRT1 (12–14).

Anaplerotic pathways are essential to maintain physiological homeostasis under carbohydrate deprivation states. Increased utilization of glutamine via α-ketoglutarate (α-KG), largely determined by the activity of glutamate dehydrogenase (GDH) within the mitochondria, is important under both normal and pathophysiological conditions, including cancer (15, 16). Interestingly, both α-KG and glutamine have been recently reported to impact TORC1 signalling (17, 18). However, under normal physiological settings if/how differential mitochondrial uptake and utilization of glutamine affect TORC1 is still unknown. In this context, mTOR is both activated and inhibited by ɑ-KG (18, 19). While glutaminolysis that generates ɑ-KG was shown to induce mTOR in cancer cells (18), ɑ-KG-mediated inhibition of mitochondrial ATP-synthase led to lifespan extension via mTOR inhibition (19). These are clearly contradictory findings and hence further investigation is required to unravel the significance of mitochondrial glutamine utilization in regulating anabolic-signalling via TORC1. Moreover, if and how mechanisms within the mitochondria that determine anaplerotic flux and energetics regulate TORC1 remains to be addressed. Specifically, it is enticing to check if mitochondrial utilization or sparing of glutamine acts as an intracellular cue to regulate metabolic signalling.

Sirtuins are typically associated with physiological responses during fasting or calorie restricted states (3). Intriguingly however, SIRT4, a *bona-fide* mitochondrial sirtuin, is induced under a fed state (20). Although, reports have shown that it has ADP-ribosyltransferase (21, 22) and NAD^+^ dependent deacylase activities (23, 24), the robust catalytic activity as well as biological functions of SIRT4 is largely unknown. More importantly, while SIRT4 is established to counter catabolic signalling and negatively regulate fatty acid oxidation (20, 21, 25), whether it affects anabolic-signalling remains unclear.

It should be noted that glutamate dehydrogenase (GDH) has been demonstrated to be a *bona-fide* substrate of SIRT4 and this is critical for glutamine homeostasis (22, 26, 27). Particularly in cancers, SIRT4-GDH-glutamine axis is known to affect cell proliferation (28) and TORC1 is known to inhibit SIRT4 expression (29). However, the link between SIRT4 and TORC1 under normal physiological conditions needs to be unravelled. It is important to note that unlike in cancers, both SIRT4 and TORC1 are induced under fed states (20). Moreover, mTOR-dependent anabolic pathways including protein synthesis and lipogenesis are energetically demanding and aberrant activation of mTOR has been shown to cause energetic stress (30). Hence, given that SIRT4 expression is higher in a fed state, it is enticing to investigate if coupling of mitochondrial metabolism/energetics with mTOR is brought about by SIRT4.

Here, we report that anaplerotic flux, specifically, utilization/sparing of glutamine in the TCA cycle constitutes a key signal mediating TORC1 activation. We, further demonstrate that SIRT4 presence or absence phenocopies TORC1 signalling under a nutrient replete or deprived state, respectively. Thus, our study provides insights into the crucial role of SIRT4 in activating TORC1 in response to nutrients, via modulation of glutamine utilization. We also demonstrate that SIRT4 enhances lipogenesis and cell proliferation, and inhibits cellular autophagy. Together we highlight SIRT4 as an anabolic sirtuin and bring to the fore the importance of mitochondrial inputs in regulating metabolic signalling.

## Results

### Mitochondrial Glutamine sparing activates TORC1

Glutamine is a potent regulator of TORC1 activity (17, 18). However, whether altered glutamine flux under physiological conditions that mimic fed or fasted states affect mTOR signalling is still unclear. Towards this, primary hepatocytes, pre-incubated in amino acid- and serum-free media, were treated with 2 mM glutamine in the presence of either low (5 mM) or high (25 mM) glucose. mTOR signalling, as assessed by phosphorylation of downstream substrate S6K (pS6K), was low in glucose-alone supplemented conditions. Addition of glutamine led to a robust increase in mTOR activity, more than in the presence of leucine and isoleucine (Fig. 1A). Interestingly, glutamine-dependent activation of mTOR signalling was highest under high glucose conditions, over and above the other treatments (Fig. 1A). In contrast to this, Leucine/Isoleucine activated mTOR signalling to similar extents under both low- and high-glucose conditions (Fig. 1A). Consistently, we found similar glucose-dependent glutamine-mediated increase in mTOR signalling even when hepatocytes were grown in complete media in the presence of amino acids and serum (Fig. 1B). It is intriguing that the effect of glutamine on pS6K under low glucose conditions was still evident in the absence of serum and amino acids and indicates a possible graded response, which is dependent upon other inputs. Nevertheless, mTOR signalling was robustly induced in high glucose, but not under low glucose conditions, irrespective of serum and other amino acids mediated effects (Fig 1A and B). Together, these suggested that glutamine effects on mTOR are potentiated in the presence of high glucose. Importantly, this led us to further investigate if differential glutamine utilization within the mitochondria, as under fed and fasted states, controls extent of mTOR signalling.

**Figure 1:**
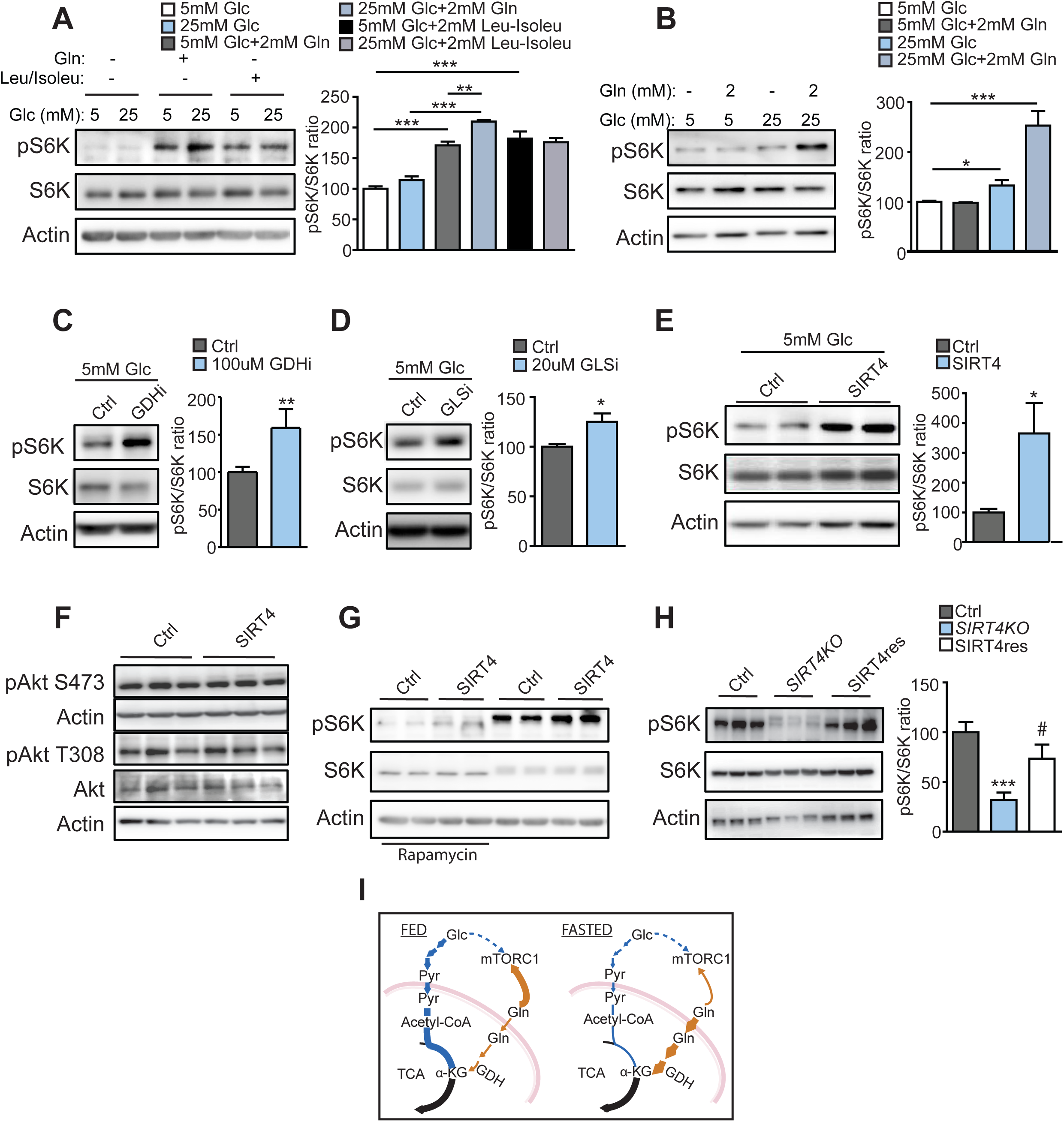
Mitochondrial SIRT4 activates cytosolic TORC1. **(A-E)** mmunoblots and quantitations for pS6K(Thr389)/S6K in **(A)** primary hepatocytes derived from wildtype mice, pre-incubated in EBSS for 3 hours, followed by 1 hour incubation in low (5 mM Glc) and high (25 mM Glc) containing EBSS with or without either 2 mM glutamine or 0.8 mM Leucine/Isoleucine (n=4); **(B)** primary hepatocytes derived from wildtype mice, under low (5 mM Glc) and high (25 mM Glc) glucose conditions with or without 2 mM glutamine (Gln) (n=5); **(C)** primary hepatocytes from wild type mice treated with 100 µM EGCG (GDHi) for 1 hour under 5 mM glucose (n=4); **(D)** primary hepatocytes from wildtype mice treated with 20μM BPTES (GLSi) for 1 hour under 5 mM glucose (n=4); **(E)** primary hepatocytes from wild type mice adenovirally transduced with Ad-CMV (control) and Ad SIRT4 (SIRT4) under low glucose conditions (n=6); **(F)** Representative immunoblots for pAkt (Ser473), pAkt (Thr308), Akt and actin in HEK293 T cells overexpressing control or SIRT4 (SIRT4-HA) under low glucose media (n=4); **(G)** Representative immunoblots for pS6K (Thr389)/S6K in primary hepatocytes adenovirally transduced with either control or SIRT4 and incubated in low glucose media with or without 20nM Rapamycin **(H)** primary hepatocytes isolated from wild type and SIRT4-KO mice. SIRT4 expression was restored by transducing Ad-SIRT4 into SIRT4-KO hepatocytes (SIRT4res) to rescue mTORC1 signaling (n=6); **(I)** Relative contributions of glucose (Glc) and glutamine (Gln) to the TCA cycle and mTORC1 activation under fed versus fasted states. pyr: pyruvate; GDH: glutamate dehydrogenase; TCA: tricarboxylic acid cycle; Data is represented as means ± SD (*p<0.05, ***p<0.001, ^#^p<0.0001).

Reduced carbohydrate availability increases anaplerotic flux of glutamine into the TCA cycle, which is regulated by the activity of GDH (31). Thus, we surmised that inhibition of GDH, which would lower glutamine channelling into TCA, could potentiate mTOR signalling. As evident from Figure 1C, a short-term inhibition of GDH in primary hepatocytes led to significant increase in pS6K. It should be noted that this increase was seen even when cells were grown under low glucose conditions, possibly due to reduced utilization of glutamine via the TCA. To confirm if this was indeed true, we also inhibited glutaminase (GLS) that converts glutamine to glutamate, which is then fed into TCA via GDH. Inhibition of GLS led to a significant increase in pS6K levels (Fig 1D) and the effects were similar to GDH inhibition (Fig 1C). It is important to note that activation of mTOR signalling following GDH or GLS inhibition under low glucose conditions was comparable to glutamine supplementation under high glucose conditions. Together these results demonstrate that, under fasted and fed conditions, differential glutamine utilization (or sparing) by the mitochondria is used as a nutrient cue to regulate mTOR in the cytosol.

### SIRT4 regulates TORC1 signalling

GDH activity and thus glutamine utilization in the mitochondria is known to be highly regulated during fed and fast conditions (31). Thus, we wanted to identify the molecular factor within the mitochondria that would mediate such effects on mTOR via glutamine. Among others, mitochondrial deacylase SIRT4 has been shown to be a potent regulator of GDH activity and anaplerotic flux (21, 22). Moreover, although SIRT4 is induced under a fed state (20, 32), functional significance of elevated SIRT4 levels in nutrient excess conditions is still unknown. Specifically, whether differential SIRT4 levels couple glutamine flux through TCA to control anabolic-signalling remains to be addressed.

To investigate this, we used either SIRT4 gain-of-function (ectopic expression) or loss-of-function (knockdown or knock out) models under different metabolic states. Given that SIRT4 levels and mTOR signalling is low under fasted conditions, we assessed the effects of SIRT4 overexpression under fasted (or low glucose) conditions. We found that ectopic expression of SIRT4 led to a robust increase in pS6K, across cell types (Figs 1E, S1A and C, S2A). mTOR is known to exist in two complexes viz. TORC1 and TORC2, and phosphorylation of pS6K and pAKT-S473 are typically used as bona fide indicators of signalling via either of these arms. On assessing pAKT-S473 under similar conditions, we did not find SIRT4-dependent TORC2 activation (Fig 1F). This highlighted that SIRT4 has a specific effect on mTORC1 signalling. Rapamycin, the well-known inhibitor of mTOR has been used to decrease signalling downstream to mTORC1, particularly at low doses (33). Consistently, we found that treating with Rapamycin significantly reduced pS6K levels in both control and SIRT4 overexpressing cells (Fig 1G). Thus together, our data demonstrates that SIRT4 positively regulates TORC1 signalling, which is sensitive to rapamycin.

Although, TORC1 has several downstream target proteins, emerging literature indicates that phosphorylation could be highly specific based on both substrate affinities and extent of activation (34, 35). Hence, we wanted to check if SIRT4-dependent activation of TORC1 led to phosphorylation of 4E-BP1, a translation repressor protein and ULK-1, which is involved in autophagy. We were surprised to find that while SIRT4-dependent TORC1 activation led to an increase in pS6K (Fig 1E, S2A) and pULK (S757) levels (Fig S2B), phosphorylation of 4E-BP1 was unaltered (Fig S2C). Although intriguing, it is now well established that 4E-BP1 is a preferred substrate of TORC1 and whereas its phosphorylation is resistant to Rapamycin treatment (33), complete starvation leads to a loss of p4E-BP1. These suggest that while minimal TORC1 activity is sufficient to phosphorylate 4E-BP1 (possibly maximally), phosphorylation of substrates like S6K is dependent on extent of mTOR activation.

Consistent with the results described above, knock down of SIRT4 led to a significant decrease in TORC1 signalling (Figs S2D-E and G and S1B). Here again, phosphorylation of 4E-BP1 remained unaltered, similar to when SIRT4 was ectopically expressed, indicating differential effect with regards to SIRT4-dependent control of TORC1 (Figs S2F). Importantly, TORC1 signalling was drastically reduced in primary hepatocytes from *SIRT4*^−/–^ mice when compared to the controls (Fig 1H). Furthermore, restoring SIRT4 expression in primary hepatocytes derived from *SIRT4*^−/–^ mice increased pS6K to levels comparable to controls, indicating rescue of TORC1 signalling (Figs 1H and S1D). These experiments using knock down or knockout of SIRT4 not only ruled out the possibility of overexpression-based artefacts, but clearly established SIRT4 as a key determinant of TORC1 signalling.

Even though until now SIRT4-dependent control of TORC1 was unknown, mTOR has been previously shown to negatively regulate *Sirt4* expression. But it should be noted that, this was shown in cancer cells (29) and it is unlikely to apply to normal physiological contexts. Importantly, SIRT4 protein levels and TORC1 signalling are highest under nutrient excess or fed states, across cells and tissues (2, 20, 25). Hence, given that both are induced in a fed state it is difficult to envisage a negative interaction between these factors under normal physiological conditions. Nonetheless, we wanted to check if mTOR inhibition affected *Sirt4* mRNA levels in primary hepatocytes. Unlike in cancer cells (29), we did not find *Sirt4* expression to be altered in response to Rapamycin (Fig S2H). These results clearly demonstrate that mitochondrial SIRT4 is a positive regulator of TORC1 signalling in non-cancerous cells/tissues.

### SIRT4 potentiates nutrient- and growth factor-dependent activation of mTORC1

Next, we wanted to explore the possibility of mitochondrial SIRT4 in potentiating or eliciting maximal TORC1 signalling. Interestingly, ectopic expression of SIRT4 under low glucose conditions, when its expression is otherwise diminished (Figs S1A and C), significantly upregulated TORC1 mediated phosphorylation of S6K, across cell types including in primary hepatocytes (Figs 2A and S3A-B). Notably, this was comparable to signalling in control-transfected cells, which were grown under high glucose conditions. Importantly, *SIRT4^−/−^* hepatocytes under both low and high glucose conditions exhibit a robust decrease in pS6K, which was rescued upon SIRT4 expression (Fig 2B), and this was reminiscent of control-transfected cells grown in low glucose containing media (Fig 2A). These results clearly indicated that while ectopic expression of SIRT4 mimics mTORC1 signalling under a high nutrient state, its knockdown phenocopied signalling in low glucose conditions.

**Figure 2:**
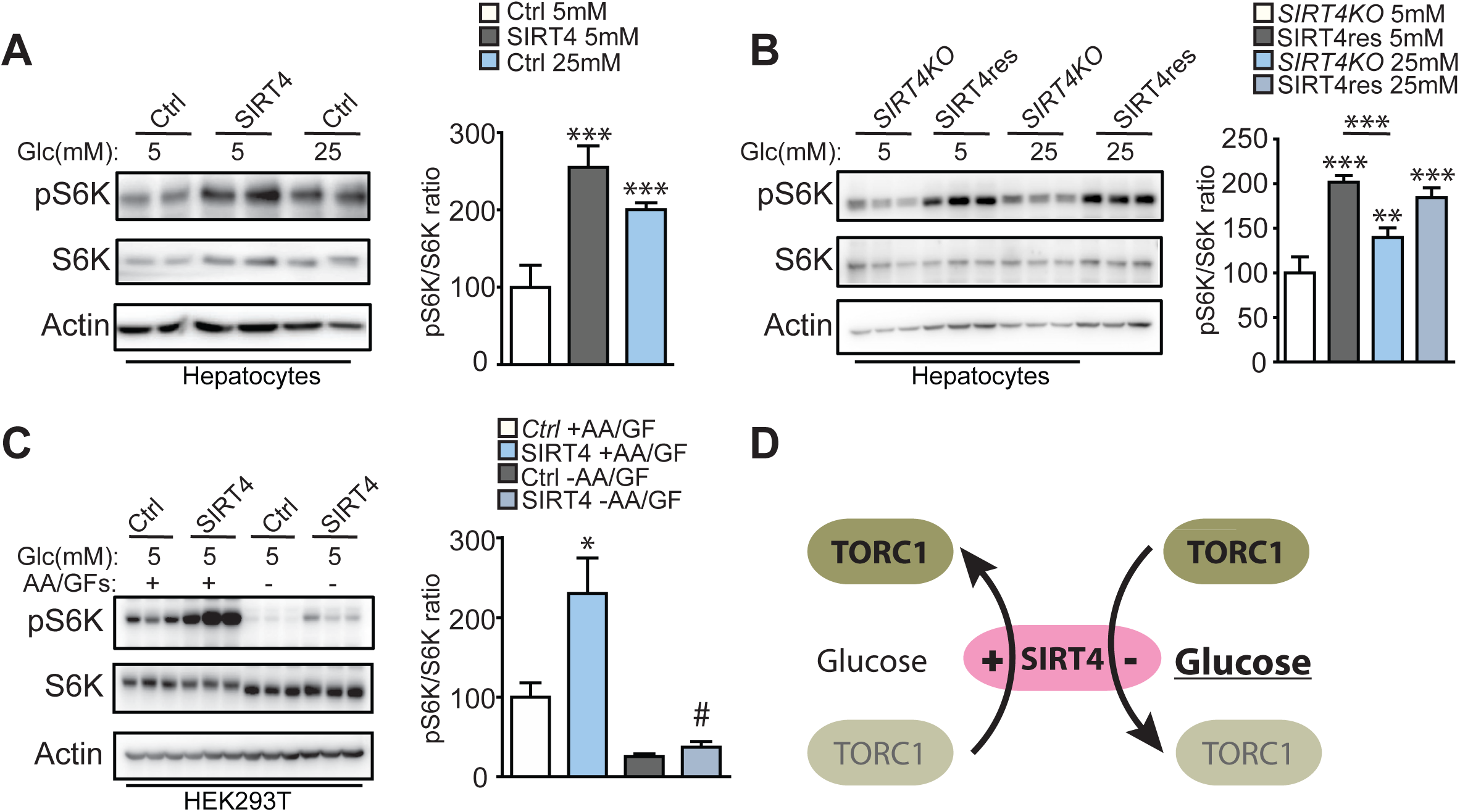
Sirt4 exerts control over nutrient and growth factor dependent activation of mTORC1. **(A)** Immunoblots and quantitations for pS6K/S6K and Actin, as indicated, in **(A)** primary hepatocytes from wild type mice transduced with Ad-CMV (control) and Ad-SIRT4 (SIRT4), and cultured in low (5 mM) and high (25 mM) glucose medium for 12 hours (n=5); **(B)** Primary hepatocytes isolated from SIRT4-KO mice. SIRT4 expression was restored by transducing Ad-SIRT4 (SIRT4) into SIRT4-KO hepatocytes (SIRT4res), and cultured in low (5 mM) and high (25 mM) glucose medium for 12 hours, as depicted (n=6); **(C)** Immunoblots for pS6K/S6K and Actin, as indicated, in control and SIRT4 transfected HEK293 T cells pre-treated with low (5 mM) glucose media for 3 hours (+AA/GFs) followed by 1 hour-EBSS treatment for serum and amino acid deprivation (-AA/GF) (n=6). **(D)** Schematic representation of the role of SIRT4 in regulating nutrient dependent activation of mTORC1. Presence of SIRT4, even under low glucose states activates mTORC1 while knockdown under high glucose states leads to attenuated mTORC1. Data is represented as means ± SD (**p<0.005, ***p<0.001, ^#^p<0.0001).

Next, given that nutrients and growth factors in the serum are potent activators of mTOR, we wanted to assess the interplay between serum/amino-acid inputs and SIRT4 in regulating TORC1 signalling. As expected, serum- and amino acid-deprivation led to a stark decrease in TORC1 signalling in both control and SIRT4 transfected cells (Fig 2C). Despite this decrease, we noticed that pS6K levels were still higher in SIRT4 transfected cells following serum- and amino acid-withdrawal, compared to control transfected cells (Figs 2C). This was interesting and indicated that ectopic expression of SIRT4 led to sustenance of TORC1 signalling even after upstream inputs were withdrawn. This could also have possibly arisen due to delayed attenuation kinetics of mTOR signalling, which needs to be investigated in the future. These results clearly indicate that levels of SIRT4 lead to differential TORC1 signalling in response to various inputs, which have been otherwise shown to be key for TORC1 activation. Importantly, SIRT4 seems to potentiate and hence, regulate anabolic-signalling mediated by TORC1.

### SIRT4-dependent glutamine sparing mediates TORC1 activation

It was striking to note the similarity between the glutamine-mediated increase in TORC1 signalling under high glucose and in a SIRT4-dependent response under low glucose (Fig 1). This prompted us to ask if SIRT4, whose expression is highest under a fed state, exerted control over TORC1 signalling via glutamine. Inhibiting mitochondrial glutamine utilization in control transfected cells, using EGCG or BPTES (inhibitors of GDH and GLS respectively), as shown earlier (Figs 1C-D), led to significant increase in pS6K levels and this increase was comparable to that observed in cells transfected with SIRT4 (Figs 3A and B). Interestingly, EGCG and BPTES treatments in SIRT4 transfected cells led to a further increase in TORC1 signalling as assessed by pS6K/S6K ratios. This strongly suggested, for most part, the likelihood of SIRT4-dependent regulation of TORC1 being brought about by differential utilization of glutamine in the mitochondria. Moreover, earlier studies both in cancers and in beta-islets have clearly established SIRT4 as a key factor in regulating glutamine utilization via the TCA cycle, which is brought about by SIRT4-mediated inhibition of GDH activity (22, 36). Thus, we hypothesized that mechanistically SIRT4-dependent glutamine sparing by the mitochondria might activate TORC1, as under a fed state.

**Figure 3:**
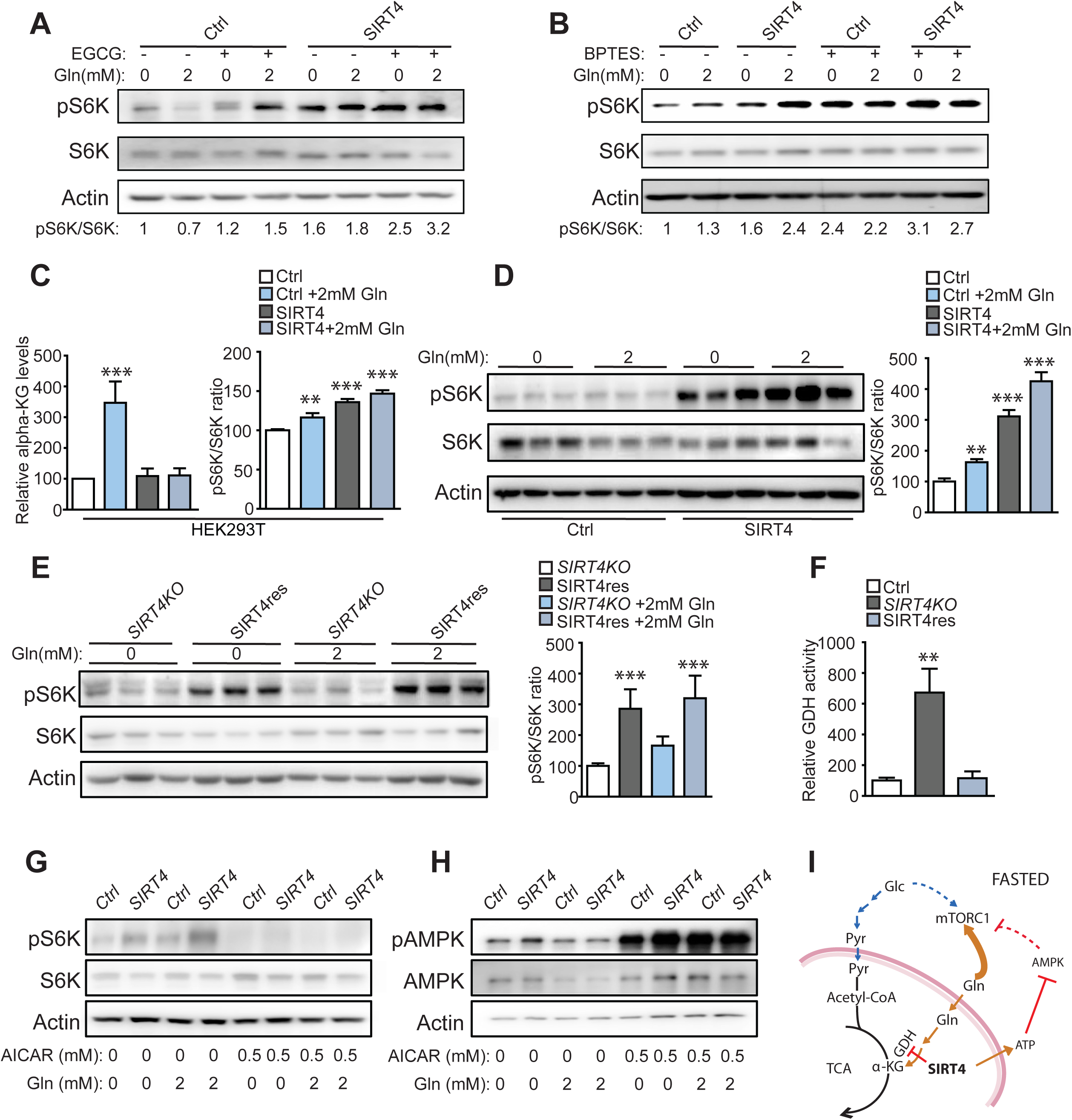
SIRT4-dependent glutamine sparing mediates TORC1 activation. **(A-B)** Representative immunoblots for pS6K/S6K and actin in primary hepatocytes derived from wildtype mice were adenovirally transduced with control or SIRT4 and incubated in low (5 mM) glucose containing media supplemented with or without 2 mM glutamine and **(A)** 100uM EGCG or **(B)** 20uM BPTES, as indicated **(C)** Relative ɑ-Ketoglutarate levels in control (Ctrl) and SIRT4 transfected (SIRT4) HEK293 T cells (n=3); and quantitations for pS6K/S6K in control (Ctrl) and SIRT4 transfected (SIRT4) HEK293 T cells (n=4) incubated in 5 mM glucose containing media supplemented with or without 2 mM glutamine; **(D-E)** Immunoblots and quantitations for pS6K/S6K and actin in primary hepatocytes derived from **(D)** wildtype (n=5) and **(E)** SIRT4-KO mice (n=6), and adenovirally transduced with Ad-CMV (Ctrl) or Ad-SIRT4 (SIRT4/SIRT4res); All experiments with glutamine were done at 2 mM glutamine (2 mM Gln) given in 5 mM glucose media for 1 hour. **(F)** Relative Glutamate dehydrogenase (GDH) activity in primary hepatocytes isolated from wildtype and SIRT4-KO mice. SIRT4 expression was restored by transducing Ad-SIRT4 in SIRT4-KO hepatocytes (SIRT4res) (n=3); **(G-H)** Representative immunoblots for **(G)** pS6K/S6K and **(H)** pAMPK/AMPK in primary hepatocytes transduced with Ad-CMV (Ctrl) or Ad-SIRT4 (SIRT4) and incubated in 5 mM glucose media with/without 2 mM glutamine and/or 0.5 mM AICAR for 1 hour (n=4); **(I)** Schematic representation of a change in anaplerotic flux driven by SIRT4, leading to mTORC1 activation even under a fasted state. SIRT4 expression, leads to increased ATP levels in the cell which lead to reduced AMPK activation and further de-repression of mTORC1 activity; Data is represented as means ± SD (**p<0.005, ***p<0.001).

Towards this, we first checked if the observed effects of SIRT4 on TORC1 were because of altered glutamine channelling into the TCA cycle, specifically under low glucose conditions. As expected, we found that glutamine supplementation led to elevated ɑ-KG levels, which was accompanied by a small increase in TORC1 signalling in control cells (Fig 3C). α-KG levels did not decrease upon SIRT4 overexpression in low glucose containing media without glutamine supplementation, nonetheless this led to activation of TORC1 (Fig 3C). However, it is important to note that unlike in control cells, while a-KG levels did not increase in SIRT4 transfected cells following glutamine supplementation, phosphorylation of S6K was significantly higher (Fig 3C). Thus, the reciprocal changes in glutamine utilization and extent of S6K phosphorylation, which was dependent upon SIRT4 expression, clearly indicated that differential channelling of glutamine into TCA cycle regulated TORC1 in the cytosol (Fig 3C and D). These findings were corroborated by enhanced GDH activity in primary hepatocytes from *SIRT4*^−/−^ mice, which was reduced to control levels when SIRT4 expression was restored (Fig 3F, S4B and C). Importantly, rescuing SIRT4 in primary hepatocytes isolated from *SIRT4*^−/−^ mice, which results in rescue of pS6K to wildtype levels (Fig 1H), led to a similar response to glutamine as is seen in hepatocytes derived from wildtype mice (Figs 3E). Based on these results, we conclude that SIRT4-mediated ‘glutamine sparing’ contributes to TORC1 activation.

### SIRT4-AMPK axis also exerts control over TORC1 signalling

Together with the results presented in Fig 3A and 3B, the additive increase in pS6K following glutamine and SIRT4 supplementation suggested that SIRT4-mediated control of TORC1 was also dependent upon another cue. We have earlier established that in the absence of SIRT4, reduced cellular ATP (and an increase in AMP/ATP ratio) leads to AMPK activation (20). Further, AMPK is an upstream inhibitor of TORC1 (37). Thus, we wanted to assess whether SIRT4-mediated inhibition of AMPK also contributed to TORC1 activation. Activation of AMPK by AICAR led to a drastic reduction in pS6K in both control and SIRT4 over expressing cells (Fig. 3G), and this decrease was not rescued by glutamine supplementation. Conversely, inhibiting AMPK in SIRT4 KO hepatocytes using compound-C led to an increase in pS6K levels. Importantly, this was further enhanced upon SIRT4 rescue (Fig. S4E). Together, these results clearly established that SIRT4 exerted a dual control over TORC1 signalling i.e. via both glutamine and AMPK.

### SIRT4 regulates mTOR localization to the lysosomes

Lysosomal localization of mTOR is essential for its activation (38) and since it is a direct readout of the extent of activation, we assessed the same as a function of SIRT4 expression in primary hepatocytes from control and *SIRT4*^−/−^ mice. We observed a drastic reduction in mTOR localization to lysosomes in *SIRT4*^−/−^ hepatocytes compared to wild type under low glucose conditions, which was rescued upon SIRT4 expression (Fig 4A, C). Importantly, there was a marked increase in mTOR intensity on the lysosomes in glutamine supplemented SIRT4 overexpressing cells (Fig 4B-C). These not only corroborated the biochemical data but also confirmed that SIRT4 was essential for activation of anabolic-signalling via TORC1.

**Figure 4:**
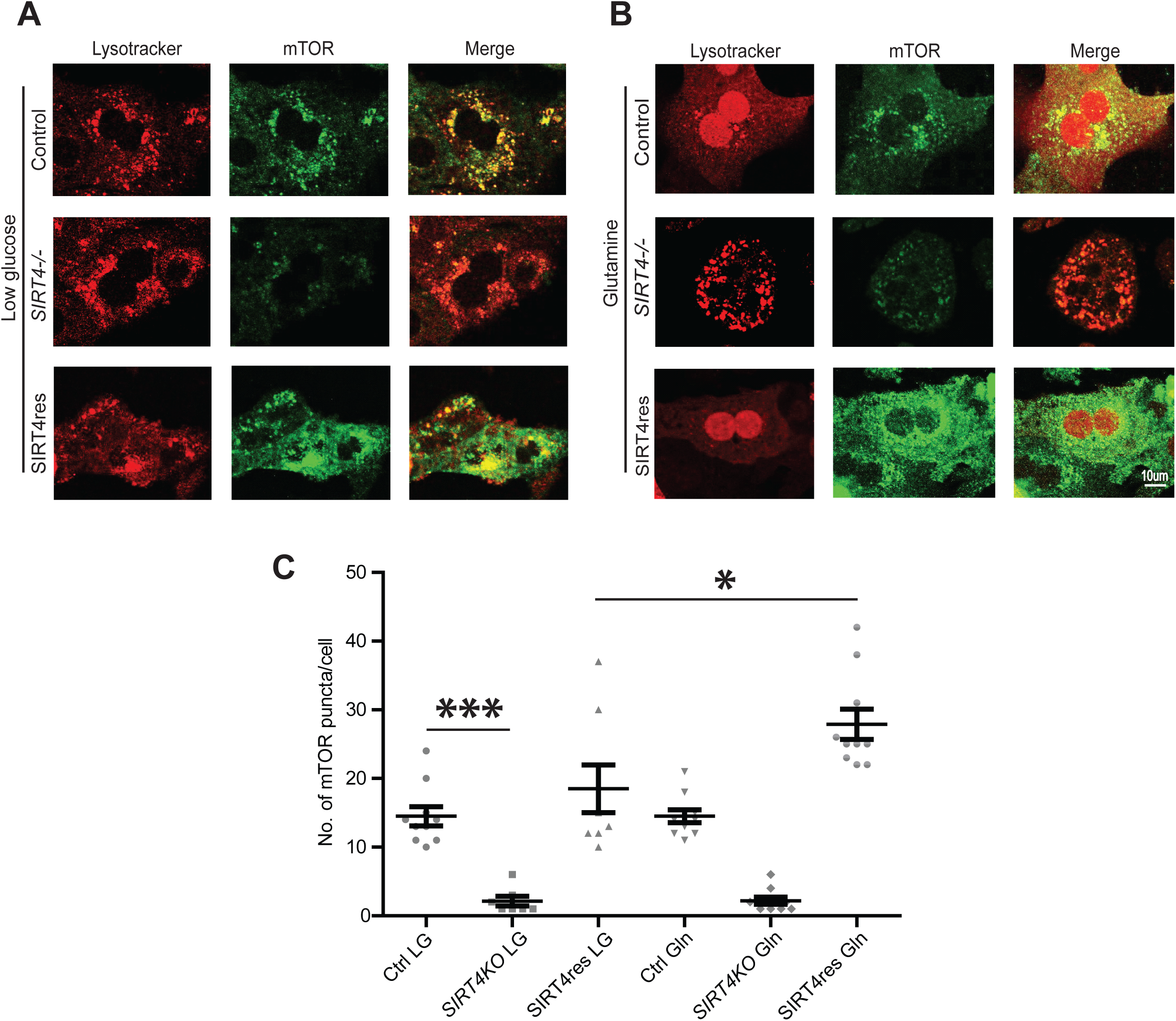
Sirt4 regulates mTORC1 localization to the lysosomes. **(A-C)** Immunofluorescence images and quantitation of mTOR puncta (green) localised to lysosomes (red) in primary hepatocytes derived from wildtype and SIRT4^−/−^ (SIRT4KO) mice, adenovirally transduced with Ad-CMV (Ctrl) or Ad-SIRT4 (SIRT4res), treated with 2 mM glutamine under low glucose (5 mM) conditions for 1 hour. Scale bar represents 40X magnification. Data is represented as means ± SEM. (*p<0.05, ***p<0.001).

### SIRT4-TOR axis impinges on transcription via SREBP1 activation

In lipogenic cells, activation of SREBP1c, the master transcriptional regulator of lipid metabolism, is primarily dependent on TORC1 activity (39). Although, previously others and we have established SIRT4 as a negative regulator of transcription of fatty acid oxidation genes, if/how it controls transcription of lipogenic genes is still unknown. Thus, we assessed if SIRT4-TORC1 axis leads to SREBP1c activation, again specifically under low glucose conditions to negate for other inputs. Ectopic expression of SIRT4 led to a robust increase in luciferase expression downstream to FAS (Fatty acid Synthetase) promoter under low glucose conditions, which was equivalent to high glucose states (Fig. 5A). Further, we found that the endogenous levels of transcripts of lipogenic genes such as FAS and SCD1 (Stearyl CoA Desaturase) were low in *SIRT4*^−/−^ hepatocytes (Fig 5B). Importantly, expression of these SREBP1c target genes were restored to wild type levels when SIRT4 expression was rescued in *SIRT4*^−/−^ hepatocytes. (Fig. 5B). Interestingly, glutamine supplementation to cells overexpressing SIRT4 led to a robust increase in FAS expression even under low glucose conditions (Fig. 5C). However, glutamine supplementation to *SIRT4*^−/−^ hepatocytes led to no increase in FAS expression, which was rescued upon SIRT4 restoration (Fig. 5D). Further, on assaying for lipid content by oil red staining, we found that *SIRT4*^−/−^ hepatocytes had reduced lipid droplets, which was again restored to control level following rescue of SIRT4 expression (Fig. S5). These clearly demonstrate that SIRT4 activates lipogenic response in hepatocytes and together with previous findings (32), indicate that it maintains the balance between lipid synthesis and breakdown.

**Figure 5:**
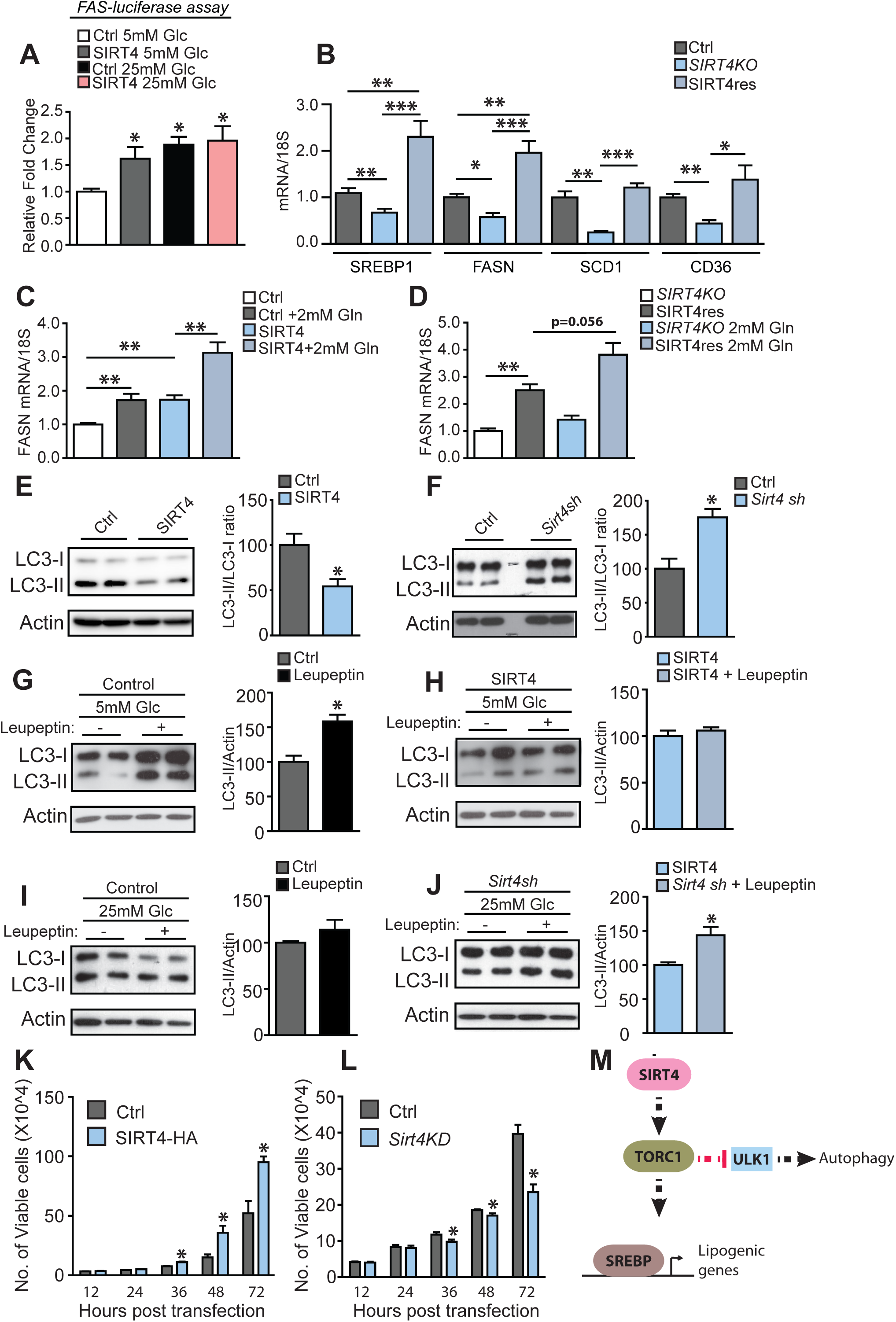
Sirt4-TOR axis impinges on transcription via SREBP1 activation. **(A)** FAS promoter driven luciferase assay in HepG2 cells over expressing SIRT4 in 5 mM and 25 mM glucose media conditions (n=6); **(B)** Quantitative RT-PCR of genes involved in lipogenesis and fatty acid uptake (SREBP1, FASN, SCD1 and CD36) from primary hepatocytes isolated from wildtype (Ctrl), SIRT4-/- (SIRT4KO) and Ad-SIRT4 transduced SIRT4KO (SIRT4res) (n=4-6); **(C-D)** Quantitative RT-PCR of FASN in primary hepatocytes derived from **(C)** wildtype and **(D)** SIRT4-/- mice, adenovirally transduced with either Ad-CMV (Ctrl/*SIRT4KO*) or Ad-SIRT4 (SIRT4/SIRT4res) and treated with 2 mM glutamine in low (5 mM) glucose media for 6 hours (n=5-6). **(E-F)** Representative immunoblots and quantitations for LC3-II/LC3-I, and actin in HEK293 T cells, transfected with **(E)** control and SIRT4 over expression vectors, kept in low (5 mM) glucose media or **(F)** control and SIRT4 knock down vectors, and kept in high (25 mM) glucose media; **(G-H)** Representative immunoblots and quantitations for LC3-II/actin in **(G)** control transfected or **(H)** SIRT4 over expressing HEK293 T cells maintained in low (5 mM) glucose media, with or without 100uM leupeptin treatment for 12 hours; **(I-J)** Representative immunoblots and quantitations for LC3-II/actin in **(I)** control knock down or **(J)** SIRT4 knockdown HEK293 T cells maintained in high (25 mM) glucose media, with or without 100uM leupeptin treatment for 12 hours; **(K-L)** Proliferation assay showing the number of cells in **(K)** HEK293 T cells transiently transfected with control and SIRT4-HA and **(L)** control and Sirt4 knock down (Sirt4KD) HEK293 T cells at different time points after plating. **(M)** Schematic representation of SIRT4-mediated control of lipogenesis and autophagy via mTORC1-SREBP1 and mTORC1-ULK1 axis respectively; Statistical significance was calculated using student’s *t-test* and ANOVA: * p < 0.05, ** p < 0.01, *** p < 0.001. Error bars indicate mean values ± SEM.

**Figure 6:**
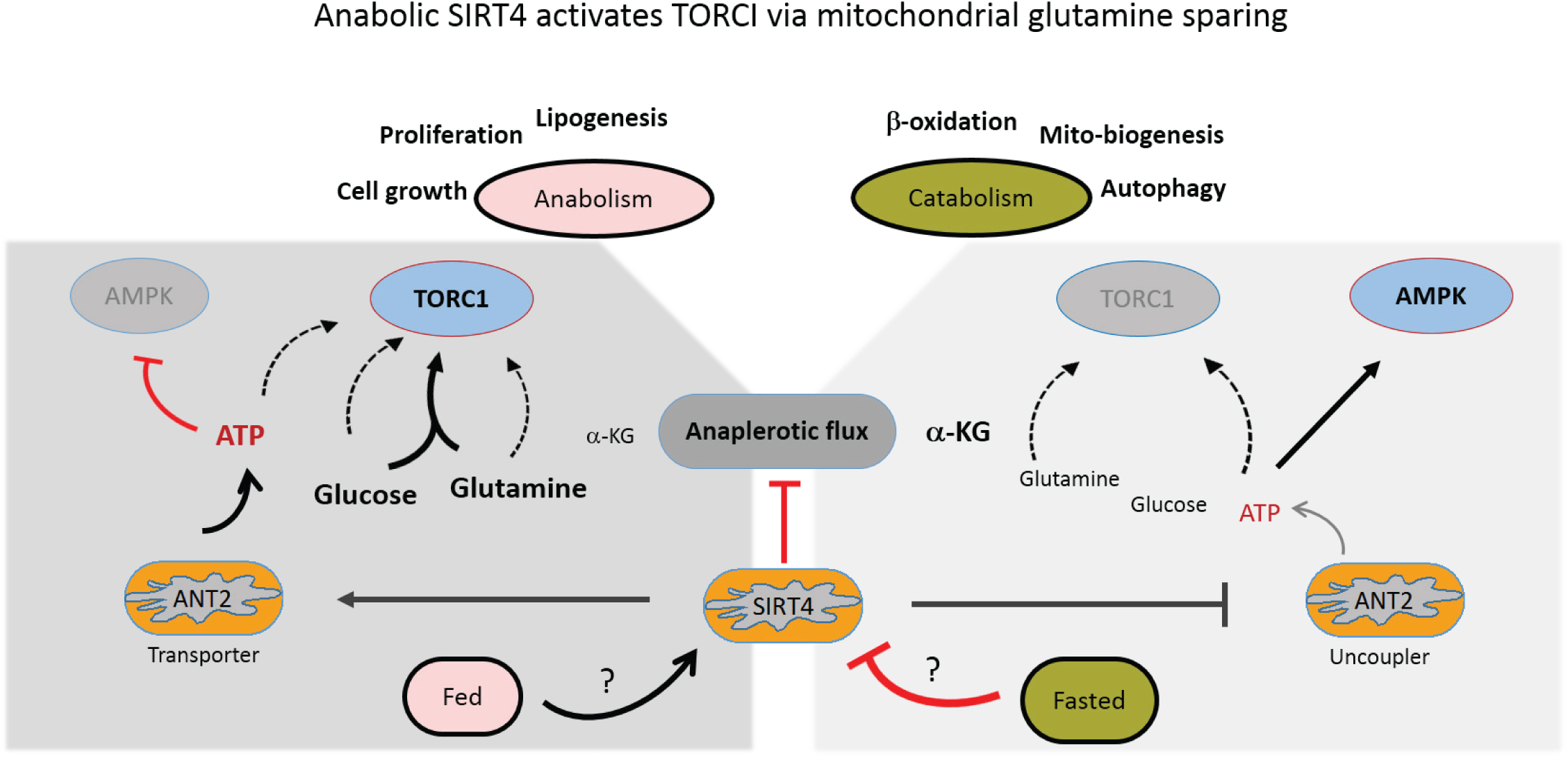
Schematic representation of mitochondrial regulation of nutrient dependent TORC1 signalling by SIRT4. SIRT4 modulates mitochondrial utilization of glutamine, which impinges on mTORC1 activation under a fasted to fed transition. Under fed states, inhibition of anaplerotic flux (via inhibition of GDH) by SIRT4, leads to spared glutamine in the cytosol, which then activates mTORC1. Under fasted states, when SIRT4 is low, increased GDH activation leads to conversion of glutamine to α-KG and hence inhibition of mTORC1 activity.

### Anabolic SIRT4 exerts control over autophagy and cell proliferation

Balance between AMPK and mTOR (specifically TORC1) signalling is critical to couple cellular metabolic/energetic state to catabolic/anabolic pathways, which determine processes like autophagy and cell proliferation/growth. Taking together our previous report on SIRT4-dependent inhibition of AMPK and the current findings on the ability of SIRT4 to activate TORC1, we wanted to assess the physiological relevance of SIRT4 expression. Specifically, we wanted to ascertain if anabolic SIRT4 could impinge on autophagy and cell proliferation.

On assessing the ratio of LC3-II to LC3-I, an indicator of autophagy, we saw a significant decrease in LC-II/LC3-I in SIRT4 over expressing cells (Fig 5E). Conversely, there was a robust increase in LC3-II/LC3-I ratio in SIRT4 knockdown cells (Fig 5F). While LC3-II/LC3-I ratio is indicative of extent of autophagy, it is not confirmatory. As established, we checked the autophagy flux by treating cells with leupeptin, which inhibits lysosomal proteases (40), and assaying for LC3-II/Actin ratio. Consistent with literature, we observed high autophagy flux in control transfected cells under low glucose conditions (Fig 5G). Interestingly, ectopic expression of SIRT4 under low glucose conditions dampened this response (Fig 5H) and was comparable to autophagy flux of control transfected cells grown in media containing high glucose (Fig 5I). On the other hand, knocking down SIRT4 expression in cells grown in high glucose media led to a significant increase in LC3-II (Fig 5J), which was similar to control transfected cells under low glucose conditions (Fig 5G). These clearly indicated that presence or absence of SIRT4 affected cellular autophagy and mimicked either a fed or fasted state, respectively and was consistent with change in TORC1 signalling (Fig 2).

In the context of autophagy, AMPK and TORC1 counteract each other by phosphorylating the common downstream substrate ULK1. Notably, TORC1 mediated inhibitory phosphorylation of ULK1 at Ser-757 (pULK757) has been shown to prevent AMPK-dependent activatory phosphorylation at Ser-317/777 (41). Thus, the effects of SIRT4 on autophagy are consistent with changes in the levels of pULK757 (Fig S2B and S2E).

To get another correlate of SIRT4-dependent effects on cellular physiology, we scored for cell proliferation, which is again inherently linked with AMPK and mTOR activities (42). As anticipated while cells expressing SIRT4 were more proliferative (Figs 5K, and S1A), knocking down SIRT4 led to reduced proliferation, as compared to respective controls (Figs 5L and S1B). This was again consistent with the ability of SIRT4 to induce anabolic-signalling with TORC1.

## Discussion

Catabolic and anabolic pathways/processes are intrinsically dependent upon the ability of cells to sense nutrient availability and have been largely studied under deprivation conditions. However, if and how intracellular utilization of one macronutrient affects the ability of the other to impinge on metabolic signalling is relatively less understood. Specifically, glutamine is utilized via anaplerotic pathways under starvation (43) and is also known to activate mTOR signalling (17, 18). Hence, whether differential glutamine utilization in the mitochondria, which is dependent upon fed or fasted state of the cell, affects cytosolic TOR is still unclear. Here, we have demonstrated that glutamine sparing by the mitochondria is key for TORC1 activation. Importantly, we establish that SIRT4, a sirtuin that is particularly abundant in the fed state, plays a crucial role in regulating TORC1 signalling through glutamine utilization in the mitochondria.

Despite several studies, SIRT4 activity and its functions have remained enigmatic. More importantly, unlike all other sirtuins, which are induced in a fasted state, SIRT4 expression is highest in a fed state (20). As emphasized earlier, SIRT4 has only been described for its role as an anti-catabolic factor (25, 32). Given this, we report its role as a mediator of anabolic-signalling via TORC1, which was hitherto unknown. In this context, we have assessed the importance of SIRT4-dependent activation of anabolic TORC1 in lipogenesis, cell proliferation and autophagy. We conclusively show that while SIRT4 enhances lipogenesis and cell proliferation, it down regulates autophagy. Others and we have established that the anti-catabolic role of SIRT4 is mediated via the inhibition of AMPK, PGC1α, SIRT1 and PPARα (20, 25), factors that are also known to affect lipogenesis, cell proliferation and autophagy. Together with our earlier study, which established SIRT4 as a negative regulator of AMPK signalling (20), we now propose SIRT4 activity in the mitochondria as a key determinant of the balance between cellular catabolic and anabolic pathways exerted via AMPK and TORC1. This is particularly evident by SIRT4-dependent change in the phosphorylation status of ULK1, a substrate of both AMPK and TORC1, in favour of TORC1 mediated modification at Ser-757. It should be noted that phosphorylation at Ser-757 is known to abrogate AMPK dependent phosphorylation of ULK1, which is activatory (41). In addition to highlighting the role of SIRT4 in AMPK-TORC1 balance, it also provides mechanistic basis to the ability of SIRT4 to regulate autophagy.

Loss of SIRT4 has been established to reduce body fat and protect from high fat induced obesity (32). While others and we have shown that inhibiting fatty acid oxidation through AMPK/PGC1α-SIRT1/PPARα axis is responsible for this phenotype (20, 25), if/how SIRT4 affects lipogenesis has not been addressed, thus far. In this context, we have found that SIRT4-TORC1 interplay regulates expression of lipogenic genes downstream to SREBP1 in primary hepatocytes and these are consistent with the lean phenotype observed in *SIRT4*^−/−^ mice (32). Motivated by our findings, it will be exciting to perturb SIRT4 expression in lipogenic tissues and study its impact on organismal physiology, in the future.

One of the key highlights of our study is the unravelling of mTOR regulation by mitochondrial glutamine sparing. Although, glutamine deprivation studies indicated that it is a crucial upstream factor that is necessary to activate mTOR (17, 18), our findings clearly show that differential glutamine utilization, via the TCA, during fed and fasted states control TORC1 signalling. Although, recent reports have indicated that α-KG could impinge on mTOR and organismal lifespan (19), α-KG has been shown to have both inhibitory and activatory effects on mTOR (18, 19). In this regard, we provide a physiologically relevant context to control of TORC1 by glutamine metabolism in the mitochondria. Notably, our results suggest that glutamine could be utilized as a metabolic signal to conditionally activate mTOR dependent anabolic pathways, which is dictated by the presence or absence of other nutrients that sustain cellular energetic needs.

Even while differential glutamine channelling into TCA affecting mTOR is rather intuitive, it becomes essential to identify a molecular factor that couples cellular energetics and glutamine metabolism. In this context, we have discovered that mitochondrial sirtuin SIRT4, whose role as a regulator of glutamine metabolism has been thoroughly established particularly in cancers (28), plays a pivotal role. We clearly demonstrate that presence or absence of SIRT4 affects glutamine metabolism via glutamate dehydrogenase and thus the differential channelling of glutamine into the TCA through α-KG determines whether or not TORC1 should be induced. Thus, we provide conclusive mechanistic support to explain both glutamine sparing-mediated and SIRT4-dependent activation of TORC1. Moreover, based on our earlier study wherein we had identified SIRT4 to be a negative regulator of AMPK and the results described here, we show that SIRT4 controls TORC1 via both AMPK and glutamine. This is interesting since glutamine utilization is intrinsically coupled to cellular energetics, as mentioned earlier (31). Therefore, it is likely that glutamine sparing and mitochondrial ATP production, which affects AMPK signalling, might synergistically bring about multiple dynamic states of mTOR/TORC1 activation.

Emerging literature indicates that both extent of TORC1 activation and differential phosphorylation of downstream substrates are crucial for encoding specificity (33, 35, 44). In this regard, it is interesting to note that SIRT4-mediated control of mTOR is limited to TORC1-dependent phosphorylation of S6K and ULK1 but not 4E-BP1. It should be noted that Kang et al reported that 4E-BP1, a high affinity substrate, is also resistant to TORC1 inhibition (33, 34), analogous to our findings. Hence, it will be exciting to address, in the future, the differential control of mTOR-dependent processes by mitochondrial signals, and the impact on cellular growth and proliferation.

Our study also distinguishes the cross-talk between metabolic inputs, SIRT4 functions and cellular physiology. Consistent with the regulation of TORC1 signalling via glutamine sparing, we have found that it is indeed differential SIRT4 expression under low or high glucose conditions, which determines the ability of cells to mount TORC1 signalling. Specifically, it should be noted that absence of SIRT4 led to a drastic reduction in TORC1 signalling even under high glucose conditions and ectopic expression of SIRT4 under low glucose conditions mimicked GDH inhibited state vis-à-vis effects on TORC1. These are also supported by changes in autophagy as a function of SIRT4 expression both under high and low glucose conditions.

In conclusion, by identifying SIRT4 as a key determinant of TORC1 signalling, we have provided molecular and physiological basis for mitochondrial control of mTOR. Being the first report to describe the dependence of anabolic-signalling on SIRT4, we believe that this will motivate further studies in unravelling the need for a sirtuin to be induced during a nutrient rich state.

## Materials and Methods

### Cell lines and primary hepatocyte culture

HEK293, HEK293T and HepG2 cell lines were obtained from ATCC and were maintained in DMEM (Sigma Cat No #D7777) medium containing 25 mM glucose with 10% fetal bovine serum (FBS) (Gibco Cat. No #16000-044) unless otherwise stated. HHL-17 cell line was a kind gift from Prof. Arvind Patel (MRC - University of Glasgow Centre for Virus Research).

Primary hepatocyte isolation: Primary hepatocytes were isolated from 3-4 months old male - wild type and *SIRT4* knockout (*SIRT4*^−/−^) mice (obtained from Jackson Laboratories, #012756), maintained under standard animal house conditions and fed with standard chow diet. Animals were sedated using thiopentone and perfused through the portal vein with HBSS solution followed by DMEM low glucose containing Collagenase A (340 μg/ml) (Sigma Aldrich Cat. no # C5138). The tissue was minced in this solution and passed through a 70 µm strainer to obtain a single cell suspension. This suspension was then centrifuged at 50 g, 4°C for 5 minutes. The pellet was washed twice and plated at desired density in DMEM high glucose with 10% FBS in Collagen Type I (Sigma Aldrich Cat. No# C3867) coated plates. Media was changed to serum free medium after 6 hours of plating. All animal studies were performed using IAEC approved protocols.

### Plasmids and constructs

Human SIRT4 cDNA was cloned into pBabe-puro. pLKO.1-eGFP scrambled shRNA was a gift from Sorab Dalal (ACTREC, India). pLKO.1 Sirt4 shRNA (TRC0000018948) was purchased from Sigma Aldrich. SIRT4 was cloned into pAdtrack CMV plasmid, which was a gift from Bert Vogelstein (Addgene plasmid # 16405).

### Adenoviral and lentiviral preparation

SIRT4 was cloned into pAdtrack CMV plasmid and adenovirus was prepared as per the protocol described by Luo et al., 2007 (45). Cells were collected post expression of GFP and were lysed using hypotonic buffer [HEPES (100 mM, pH 7.5), MgCl_2_ (1.5 mM), KCl (10 mM), DTT (0.5 mM)]. Supernatant was collected and used for transduction in different cell types. SIRT4 expression was confirmed by RT-PCR/-qPCR.

For lentiviral preparation, pLKO.1 scrambled shRNA or *Sirt4* shRNA was transfected in HEK293 T cells with packaging plasmids pMD2.G (Addgene plasmid # 12259) and psPAX2 (Addgene plasmid # 12260), which were a gift from Didier Trono. Media was collected at 36 hours and 48 hours post transfection, filtered using 0.45 µm filters and stored at −80°C until use.

### Transfection and Transductions

Cells were transfected with plasmids as indicated using Lipofectamine 2000 (Invitrogen Cat no # 11668019) according to manufacturer’s instructions. Adenoviral particles were used for transducing cell lines after 12 hours of plating; cells were collected 36 hours post transduction after respective treatments. Primary hepatocyte cultures were transduced with adenovirus after 24 hours of plating and collected at 72 hours post plating after respective treatments. Lentiviral particles were used for knockdown in cells after 24 hours of plating, in the presence of polybrene (Sigma Aldrich Cat # H9268) at a concentration of 8 µg/ml. Cells were collected 48 hours post transduction.

### Treatments

All treatments were given in FBS containing media unless otherwise specified. For GDH inhibition experiments, Epigallocatechin gallate (EGCG, Sigma Aldrich Cat. No# E4143) was used at a concentration of 100uM for 1 hour in low (5 mM) glucose media. For GLS inhibition experiments, Bis-2-(5-phenylacetamido-1,3,4-thiadiazol-2-yl)ethyl sulfide (BPTES, Sigma Aldrich Cat. No# SML0601) was used at a concentration of 20μM for 1 hour in low (5 mM) glucose media. For experiments under low/high glucose, cells were either shifted to 5 mM glucose containing media (Low) (Sigma Cat No #D5523) or fresh 25 mM glucose containing media (High), 12 hours prior to collection. For TOR inhibition, cells were treated with 20 nM Rapamycin (Sigma Cat No# 0395) for 30 minutes under low glucose conditions. For amino acid and growth factor starvation, cells were kept in 5 mM glucose containing DMEM medium for 3 hours followed by 1 hour in serum free EBSS (Cat. No # E2888, Sigma) prior to collection. For glutamine supplementation experiments, cells were kept in 2 mM glutamine supplemented in 5 mM glucose or 25 mM glucose containing DMEM medium (as indicated in figure) for 1 hour. For GDH assay, cells were cultured in 5 mM glucose containing DMEM medium. For inhibition of autophagy flux, cells were kept in either low (5 mM) or high (25 mM) glucose media with or without 100μM Leupeptin (Sigma Cat No# L2884) for 12 hours. For qPCR and luciferase assays, cells were kept in the indicated media conditions for 6 hours.

### Western blotting

Cells were lysed with RIPA lysis buffer [Tris(50 mM, pH8), Sodium chloride (150 mM), SDS (0.1%), sodium deoxycholate (0.5%), triton X-100 (1%), 1 mM sucrose)] supplemented with a protease inhibitor cocktail (Roche, Catalog No: 04693159001), phosphatase inhibitor PhosSTOP (Roche, Catalog No: 000000010837091001) and 1 mM PMSF (Sigma). The lysates were centrifuged at 12,000 rpm (4°C) for 10 minutes to remove cell debris. The concentration of protein was measured using the BCA Protein Assay kit (Thermo Fisher Scientific Cat. No# 23225). Subsequently, equal amounts of protein (in 1X Laemmli loading buffer) were resolved using SDS-PAGE gels and transferred to PVDF membranes (Millipore Cat. No# IPVH00010). After blocking in 5% BSA or 5% fat-free milk in TBST (TBS with 0.1% Tween-20) for 1 hour at RT, membranes were incubated with primary antibody at 4°C overnight. To visualize the bands, blots were incubated with HRP conjugated secondary antibodies in 5% fat-free milk in TBST for 1 hour at RT followed by washes in TBST. Next, the membranes were visualized using the GE AI600 chemiluminescence system with ECL reagent from Thermo Scientific (Cat. No#1859023/185022).

### Immunofluorescence

After respective treatments and at indicated time points, primary hepatocytes plated on collagen coated cover slips, were treated with 75nM LysoTracker™ Deep Red (Thermo Fisher Scientific Cat. No# L12492) for 15 minutes in the same media. Cells were then rinsed once with PBS and fixed in chilled 4% PFA for 30 minutes. After fixation, the cells were washed thrice with PBST (PBS with 0.1% Tween-20). After blocking and permeabilization in 5% BSA and 0.5% Triton X in PBST for 40 minutes, cells were incubated overnight at 4°C, with mTOR antibody (Cat. No# 2983). Cells were then incubated with anti-rabbit Alexa Fluor 647 (Thermo Fisher Scientific Cat. No# L12492) for 1 hour at RT followed by washes in PBST. Coverslips were then mounted on a slide and imaged at 40X using FluoView® FV1200 Laser Scanning Confocal Microscope from Olympus Life Science.

### Oil Red O staining

Primary hepatocytes plated in collagen coated plates, after respective treatments, were rinsed once with PBS and fixed in chilled 4% PFA for 30 minutes. The cells were rinsed once again in PBS and freshly prepared and filtered Oil Red O solution (40% in distilled water from a 0.5% stock solution in isopropanol) was added and kept for 15 minutes. The cells were washed in distilled water. The cells were then kept in water and imaged on EVOS FLc microscope from Life technologies Inc.

### Antibodies

The following antibodies were used for Western blot analyses. Anti-phospho S6K (Thr 389) (Cat. No# 9234S), anti-p70-S6K (Cat. No# 2708S), anti-phospho ULK1 (Ser757) (Cat. no# 6888S), anti-ULK1 (Cat. No# 8054S), anti-phospho 4E-BP1 (Cat. no# 9456S), anti-4E-BP1 (Cat. no# 9644S) and anti-LC3A/B (Cat. No# 12741S) were obtained from Cell Signaling Technologies (USA). Anti-β-Actin (Cat. No # A1978), anti-HA tag (Cat. No# H6908, Sigma), anti-Rabbit secondary (Cat. No# A0545) and anti-mouse secondary (Cat. No# A9044) antibodies were obtained from Sigma Aldrich (USA).

### RNA isolation and quantitation

Total RNA was extracted using the Trizol reagent (Cat. No# 15596018) according to the manufacturer’s instructions. 1μg of RNA was reverse transcribed into cDNA using random hexamers and SuperScript IV reverse transcriptase kit (Cat. No# 18090200). PCR was carried out with KAPA SYBR® FAST Universal 2X qPCR Master Mix (Cat# KK4601) / LightCycler 480 SYBR Green I Master kit (Roche Cat# 14712220) in Vapo.Protect Eppendorf LC-480 and LC-96 from Roche system using primer pairs mentioned in Supplementary Table A.

### ɑ-Ketoglutarate (ɑ-KG) assay and Glutamate dehydrogenase (GDH) assay

ɑ-KG levels and GDH activity were assayed in cell lysates using ɑ-KG measurement kit (Cat. No# ab83431) and GDH activity kit (Cat. No# ab102527) from Abcam as per manufacturer’s instructions. Briefly, cells were lysed using assay buffer provided in the kit and centrifuged at 13000 rpm for 3-5 minutes at 4°C. For ɑ-KG, lysates were further deproteinized using perchloric acid and then processed as per manufacturer’s instructions. Both assays were set up in 96-well plates and colorimetric readings taken using Tecan Infinite M200 pro plate reader system at 570 nm for ɑ-KG and 450 nm for GDH. ɑ-KG levels and GDH activity were normalized to protein concentration (estimated using BCA: Pierce, Thermofischer Cat no# 23225).

### Luciferase assay

HepG2 cells were transfected with the luciferase expression construct under the Fatty Acid Synthetase (FAS) Promoter (Addgene# 8890) along with β-Gal plasmid (Ambion Cat. No# 5791). After 12 hours of transfection, cells were transduced with either Ad-CMV or Ad-SIRT4 for another 24 hours, following which the cells were kept for 6 hours in either low (5 mM) or high (25 mM) glucose. Cells were then harvested and lysed in passive lysis buffer (Promega Cat. No# E1941). β-galactosidase assay was performed using ONPG (orthoNitrophenyl-β-galactoside, Sigma Cat. No# N1127) as a substrate. Luciferase assay was performed with the Luciferase assay system (Promega Cat. No# E1500) as per manufacturer’s instructions. Luminescence counts were measured in Infinite-M200-Pro (Tecan) or TriStarLB941 (Berthold technologies) and normalized to the β-galactosidase values to determine relative luciferase units (RLU).

### Data processing and statistical analysis

Western blots and immunofluorescence data were analysed using ImageJ software. Student’s T-test and ANOVA were used for statistical analysis (p value: *< 0.05; ** < 0.01; *** < 0.001 or as indicated). Microsoft Excel was used for data processing, and statistical significance was calculated using Excel or GraphPad Prism 5.0. Results are given as the means ± standard deviation or as indicated. All experiments were performed at least twice with a minimum of 3-6 replicates.

## Acknowledgements

We thank Prof. Arvind Patel (MRC - University of Glasgow Centre for Virus Research) for gifting us HHL-17 cells. We thank Dr Sorab Dalal for gifting us the pLKO.1 eGFP vector. We thank Dr. Kalidas Kohale and Dr. Shital Suryavanshi (TIFR-AH) and Dr. Sachin, Dr. Sagar Tarte and Ms. Ritika Gupta (National Facility for Gene Function in Health and Disease, IISER Pune) for help with the animal experiments. We thank Himani Narang, G. Abhinav and Abhrajyoti Chakrabarti for their help with the proliferation assays.

## Author contributions

UK designed and supervised the research; ES, MT, NM and AA performed the research; and UK, ES, and MT analysed the data and wrote the manuscript.

